# Deciphering the Transcription Factor Landscape in Neuroendocrine Prostate Cancer Progression: A Novel Approach to Understand NE Transdifferentiation

**DOI:** 10.1101/2024.04.27.591428

**Authors:** Yu Wang, Hui Xue, Xiaohui Zhu, Dong Lin, Xin Dong, Zheng Chen, Junru Chen, Mingchen Shi, Yuchao Ni, Jonathan Cao, Rebecca Wu, Ning Kang, Xinyao Pang, Francesco Crea, Yen-Yi Lin, Colin C. Collins, Martin E. Gleave, Abhijit Parolia, Arul Chinnaiyan, Christopher J. Ong, Yuzhuo Wang

## Abstract

**Background and Objective:** Prostate cancer (PCa) is a leading cause of cancer mortality in men, with neuroendocrine prostate cancer (NEPC) representing a particularly resistant subtype. The role of transcription factors (TFs) in the progression from prostatic adenocarcinoma (PRAD) to NEPC is poorly understood. This study aims to identify and analyze lineage-specific TF profiles in PRAD and NEPC and illustrate their dynamic shifts during NE transdifferentiation.

**Methods:** A novel algorithmic approach was developed to evaluate the weighted expression of TFs within patient samples, enabling a nuanced understanding of TF landscapes in PCa progression and TF dynamic shifts during NE transdifferentiation.

**Results:** unveiled TF profiles for PRAD and NEPC, identifying 126 shared TFs, 46 adenocarcinoma-TFs, and 56 NEPC-TFs. Enrichment analysis across multiple clinical cohorts confirmed the lineage specificity and clinical relevance of these lineage-TFs signatures. Functional analysis revealed that lineage-TFs are implicated in pathways critical to cell development, differentiation, and lineage determination. Novel lineage-TF candidates were identified, offering potential targets for therapeutic intervention. Furthermore, our longitudinal study on NE transdifferentiation highlighted dynamic TF expression shifts and delineated a three-phase hypothesis for the process comprised of de-differentiation, dormancy, and re-differentiation. and proposing novel insights into the mechanisms of PCa progression.

**Conclusion:** The lineage-specific TF profiles in PRAD and NEPC reveal a dynamic shift in the TF landscape during PCa progression, highlighting three distinct phases of NE transdifferentiation.

## Introduction

Prostate cancer (PCa) remains a significant public health challenge, being the most commonly diagnosed cancer and the second leading cause of cancer-related mortality among men in the United States (*1*). Prostatic carcinogenesis and PCa progression are closely linked to the androgen receptor (AR) signaling pathway, which is targeted by treatments such as androgen deprivation therapy (ADT) and AR pathway inhibitors (ARPIs), including next-generation ARPIs like Enzalutamide (*2*), Apalutamide (*3*), Darolutamide (*4*) and Abiraterone (*5, 6*). Even though these therapies improve patient outcomes, their use has led to the emergence of a lethal, ARPI-resistant subtype of prostate cancer known as treatment-induced neuroendocrine prostate cancer (NEPC), accounting for 10-17% of cases after ARPI therapy (*7–9*).

The challenge presented by NEPC, underscored by its resistance to current treatments, brings to light the critical need for a deeper understanding of its development from prostatic adenocarcinoma (PRAD). Insights into the cellular plasticity of PCa, particularly regarding NEPC development, highlight that NEPC may arise from adenocarcinoma through a process known as NE transdifferentiation (*10, 11*). This NE transdifferentiation represents a pivotal lineage transition, enabling tumor cells to evade the constraints of ADT and emerge as an androgen-independent growth pattern, thereby fostering resistance to therapeutic interventions (*12*).

Whilst adenocarcinoma and NEPC may share certain genetic alterations (*13*), their transcriptomic landscapes diverge significantly, largely under the influence of their distinctive transcription factors (TFs). Studies across various biological systems have demonstrated that TF expression profiles exhibit lineage-specific restrictions during cellular differentiation, suggesting that key combinations and interactions among TFs dictate the trajectory of lineage commitment and cellular identity (*14*). In PCa, this suggests that specialized TF networks are responsible for the gene expression variances observed between adenocarcinoma and NEPC.

Emerging evidence points to the dysregulation of TFs as key oncogenic drivers in PCa progression (*15, 16*). Notably, the AR stands as a central TF in prostatic adenocarcinoma, orchestrating the expression of genes critical for disease pathogenesis (*17*). Additionally, aberrant expression and activity of TFs from the ETS family, such as ERG and ETV1, has been linked to prostatic adenocarcinoma development (*18, 19*). Moreover, TFs like FOXA1, NKX3.1, and MYC have emerged as key players in prostatic adenocarcinoma (*20–22*). Similarly, TFs such as ASCL1, BRN2, FOXA2, and ONECUT2 (*23–26*) are pivotal in NEPC, further underscoring the importance of TFs in disease biology. Despite these advancements, of the total 1639 TFs encoded by the human genome (*27*), only a minority have been associated with PRAD and NEPC. Hence, a comprehensive study on the TF landscape associated with PCa is greatly warranted. Many TFs, potentially crucial in PCa progression and NE transdifferentiation, remain unidentified or inadequately studied due to their low abundance and subtle expression changes, which might indeed regulate the expression program of those “high jumper” genes (28). This presents a significant knowledge gap, particularly in understanding TFs relevant to drug resistance and lineage transition in PCa. The gap can partly be attributed to the predominant focus on a limited number of TFs, while many others remain undiscovered or underexplored in PCa research.

To address these critical gaps, our study introduces a methodological approach designed for the comprehensive identification and analysis of lineage-specific TFs in PCa. We applied this innovative approach with the aim of elucidating the temporal dynamics and functional roles of TFs during NE transdifferentiation. We anticipate that our research will not only deepen the understanding of TF-mediated lineage plasticity and drug resistance in PCa but also unveil novel targets for therapeutic intervention, ultimately contributing to the advancement of personalized cancer treatment strategies and improving patient outcomes.

## Results

### A novel method reveals TF profiles in PRAD and NEPC

We have developed a novel algorithm to comprehensively identify PCa-related TFs, revealing distinct TF profiles for PRAD and NEPC (**Fig. 1A**). Unlike traditional methods that rely primarily on fold change—limited by its inability to reflect low-expressed, yet significantly altered genes which restricts the comprehensive representation of gene significance—our approach uses the internal Z score. This new metric assesses a TF’s weighted expression within a sample, enhancing the identification of robust lineage signatures beyond mere fold changes, thereby increasing the precision of our analysis to identify meaningful lineage signatures.

**Fig. 1.**
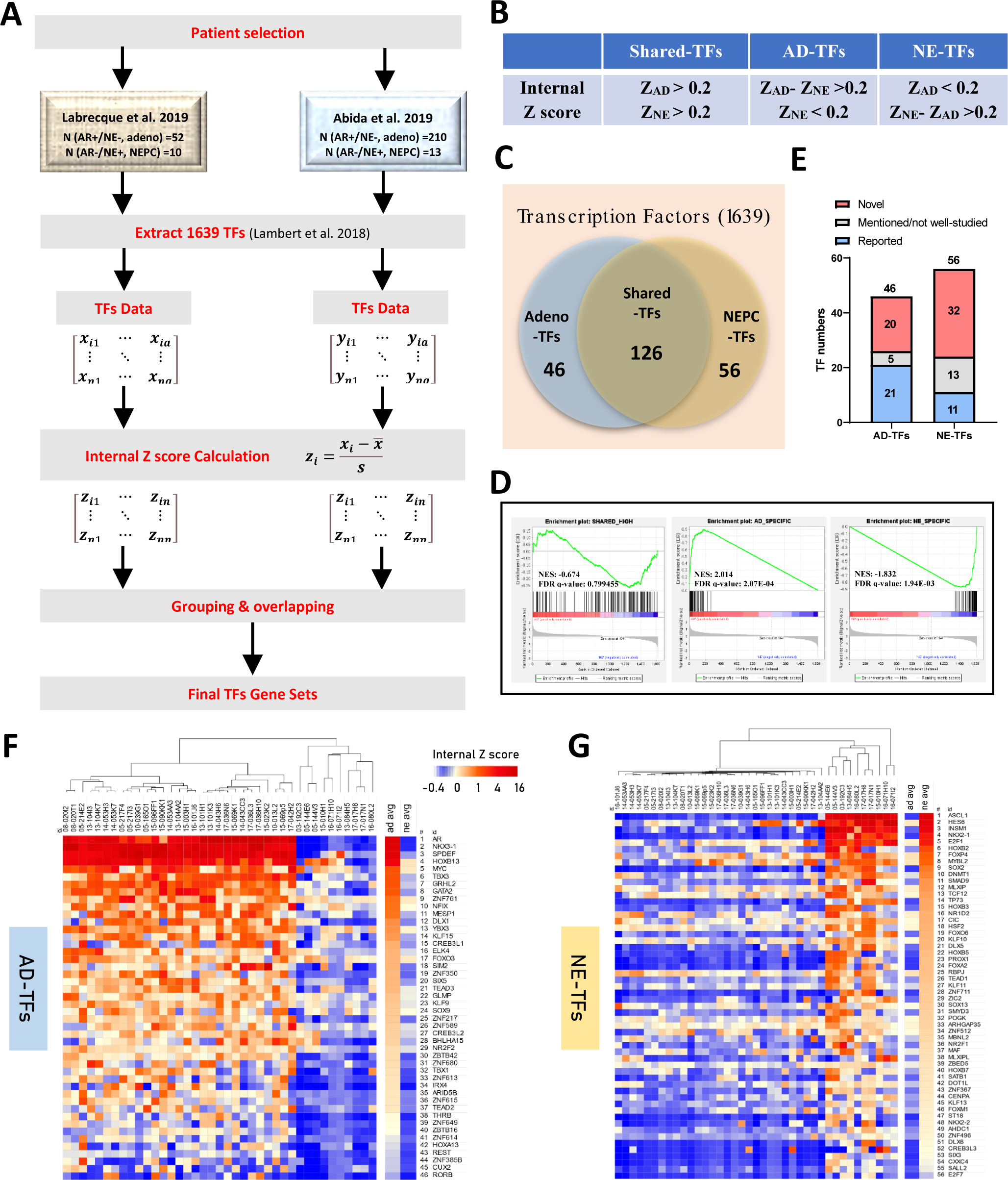
Illustration of the Lineage-TF Signatures Identification Process. (**A**) Schematic depiction of the novel algorithm developed to discern lineage-TF signatures. Initial TF expression data matrices were extracted from two distinct clinical cohorts following patient sample selection criteria. Subsequently, internal Z scores were computed to depict the weighted TF expression within each sample. Employing specified criteria, TF gene sets were derived, with final TF lists determined through overlap analysis. (**B**) Criteria for grouping. Z_AD_: the trimmed mean of internal Z score for a TF among PRAD samples. Z_NE_: the trimmed mean of internal Z score for a TF among NEPC samples. (**C**) Summary graph presenting the TF signature identification outcomes. Out of 1639 TFs examined, 126 were identified as shared-TFs, 46 as AD-TFs, and 56 as NE-TFs. (**D**) Gene set enrichment analysis in the Labrecque et al. 2019 cohort indicated enrichment of the shared-TFs gene set in both PRAD and NEPC patients (NES=-0.674, FDR q-value=0.79945). The AD-TFs gene set was enriched solely in PRAD patients (NES=2.014, FDR q-value=2.07×10−4), while the NE-TFs gene set showed exclusive enrichment in NEPC patients (NES=-1.832, FDR q-value=1.94×10−3). (**E**) A comprehensive literature review disclosed that among the 46 AD-TFs, 21 were previously identified as PRAD-related regulators, with 20 being novel discoveries. Among the 56 NE-TFs, 11 were established players in NEPC, 32 were previously unreported, and 13 had been briefly mentioned but not extensively studied. (**F-G**) Heatmaps exhibiting the signature of (**F**) AD-TFs and (**G**) NE-TFs, with hierarchical clustering revealing distinct separation between PRAD and NEPC patients, ranking TFs based on the trimmed mean of internal Z scores within respective lineages. *NES*: *Normalized Enrichment Score. FDR: False Discovery Rate*.

To minimize the noise brought by the heterogeneity of PCa, we pre-selected patient samples from the cohort, remaining the samples characterized as AR+/NE-adenocarcinoma and AR-/NE+ small cell neuroendocrine carcinoma based on previous reported specific gene panels **(Fig. S1**) (29). By applying our innovative methodology to assess the internal Z scores of 1,639 documented TFs across these samples, we effectively mapped distinct TF profiles, clarifying lineage affiliations (27). Based on our criteria (**Fig. 1B**), we classified TFs into three groups (**Fig. 1C**): 126 shared TFs with high internal Z scores across both lineages, 46 adenocarcinoma-specific TFs (AD-TFs) with high scores in adenocarcinoma but lower in NEPC (**Table 1**), and 56 NEPC-specific TFs (NE-TFs) with distinct expression in NEPC (**Table 2**).

**Table 1.** AD-TFs list.

| Rank | Gene Name | Z<sub>AD</sub> | Z<sub>NE</sub> | Z<sub>AD-ZNE</sub> | p-value | Novelty in PRAD |
| --- | --- | --- | --- | --- | --- | --- |
| 1 | AR | 12.2120 | -0.3449 | 12.5570 | 1.95E-08 | reported |
| 2 | NKX3-1 | 11.3254 | -0.3374 | 11.6628 | 1.08E-13 | reported |
| 3 | SPDEF | 10.3354 | -0.1559 | 10.4913 | 1.72E-15 | reported |
| 4 | HOXB13 | 5.7601 | 0.1648 | 5.5953 | 5.12E-11 | reported |
| 5 | MYC | 1.5457 | 0.1358 | 1.4099 | 6.30E-05 | reported |
| 6 | TBX3 | 1.5455 | -0.1175 | 1.6630 | 2.95E-10 | Not-well studied |
| 7 | GRHL2 | 1.4861 | -0.0843 | 1.5704 | 7.38E-11 | reported |
| 8 | GATA2 | 1.3587 | -0.1617 | 1.5203 | 2.90E-06 | reported |
| 9 | ZNF761 | 0.9096 | 0.1428 | 0.7668 | 6.63E-04 | Novel |
| 10 | NFIX | 0.9035 | 0.0221 | 0.8814 | 1.73E-05 | Not-well studied |
| 11 | MESP1 | 0.8833 | -0.1193 | 1.0026 | 9.54E-07 | Novel |
| 12 | DLX1 | 0.7125 | -0.1923 | 0.9048 | 1.91E-07 | reported |
| 13 | YBX3 | 0.6914 | 0.0706 | 0.6209 | 2.78E-05 | Novel |
| 14 | KLF15 | 0.6472 | -0.1362 | 0.7833 | 1.13E-06 | Novel |
| 15 | CREB3L1 | 0.5059 | -0.1678 | 0.6737 | 1.25E-04 | Not-well studied |
| 16 | ELK4 | 0.4767 | -0.0564 | 0.5331 | 2.25E-04 | reported |
| 17 | FOXO3 | 0.4733 | 0.1399 | 0.3334 | 6.91E-03 | reported |
| 18 | SIM2 | 0.4712 | -0.2193 | 0.6905 | 3.82E-02 | reported |
| 19 | ZNF350 | 0.4459 | -0.2428 | 0.6887 | 9.00E-07 | Novel |
| 20 | SIX5 | 0.4451 | -0.1577 | 0.6028 | 9.88E-07 | Novel |
| 21 | TEAD3 | 0.4406 | -0.0285 | 0.4690 | 5.11E-04 | Novel |
| 22 | GLMP | 0.3787 | -0.0229 | 0.4016 | 1.30E-05 | Novel |
| 23 | KLF9 | 0.3513 | -0.0372 | 0.3885 | 1.34E-06 | reported |
| 24 | SOX9 | 0.3342 | 0.0031 | 0.3312 | 3.87E-03 | reported |
| 25 | ZNF217 | 0.3327 | -0.2333 | 0.5660 | 3.43E-11 | Not-well studied |
| 26 | ZNF589 | 0.2979 | -0.0467 | 0.3446 | 4.08E-03 | Novel |
| 27 | CREB3L2 | 0.2933 | -0.0669 | 0.3603 | 4.18E-03 | reported |
| 28 | BHLHA15 | 0.2814 | -0.1272 | 0.4086 | 1.78E-02 | Novel |
| 29 | NR2F2 | 0.2180 | -0.0508 | 0.2689 | 6.79E-05 | reported |
| 30 | ZBTB42 | 0.2168 | -0.2798 | 0.4966 | 5.08E-11 | Novel |
| 31 | ZNF680 | 0.2133 | -0.2197 | 0.4330 | 5.44E-04 | Novel |
| 32 | TBX1 | 0.1840 | -0.2885 | 0.4725 | 5.35E-03 | reported |
| 33 | ZNF613 | 0.1736 | -0.2936 | 0.4672 | 1.08E-06 | Novel |
| 34 | IRX4 | 0.1684 | -0.3740 | 0.5424 | 6.12E-05 | reported |
| 35 | ARID5B | 0.1583 | -0.1922 | 0.3505 | 4.21E-06 | reported |
| 36 | ZNF615 | 0.1546 | -0.2037 | 0.3584 | 3.36E-05 | Novel |
| 37 | TEAD2 | 0.1516 | -0.1969 | 0.3485 | 6.83E-04 | Novel |
| 38 | THRB | 0.0307 | -0.3466 | 0.3773 | 1.93E-07 | Not-well studied |
| 39 | ZNF649 | 0.0108 | -0.2812 | 0.2920 | 7.00E-07 | Novel |
| 40 | ZBTB16 | -0.0013 | -0.2894 | 0.2881 | 2.08E-08 | reported |
| 41 | ZNF614 | -0.0351 | -0.2430 | 0.2080 | 1.39E-05 | Novel |
| 42 | HOXA13 | -0.0556 | -0.2919 | 0.2363 | 3.07E-03 | reported |
| 43 | REST | -0.0727 | -0.3126 | 0.2399 | 3.14E-08 | reported |
| 44 | ZNF385B | -0.0812 | -0.2986 | 0.2173 | 6.56E-03 | Novel |
| 45 | CUX2 | -0.0836 | -0.2843 | 0.2007 | 4.46E-03 | Novel |
| 46 | RORB | -0.1338 | -0.3667 | 0.2329 | 2.31E-03 | Novel |

Z<sub>AD</sub>: Trimmed mean of internal Z score among prostatic adenocarcinoma patient samples.Z<sub>NE</sub>: Trimmed mean of internal Z score among NEPC patient samples.

**Table 2.**
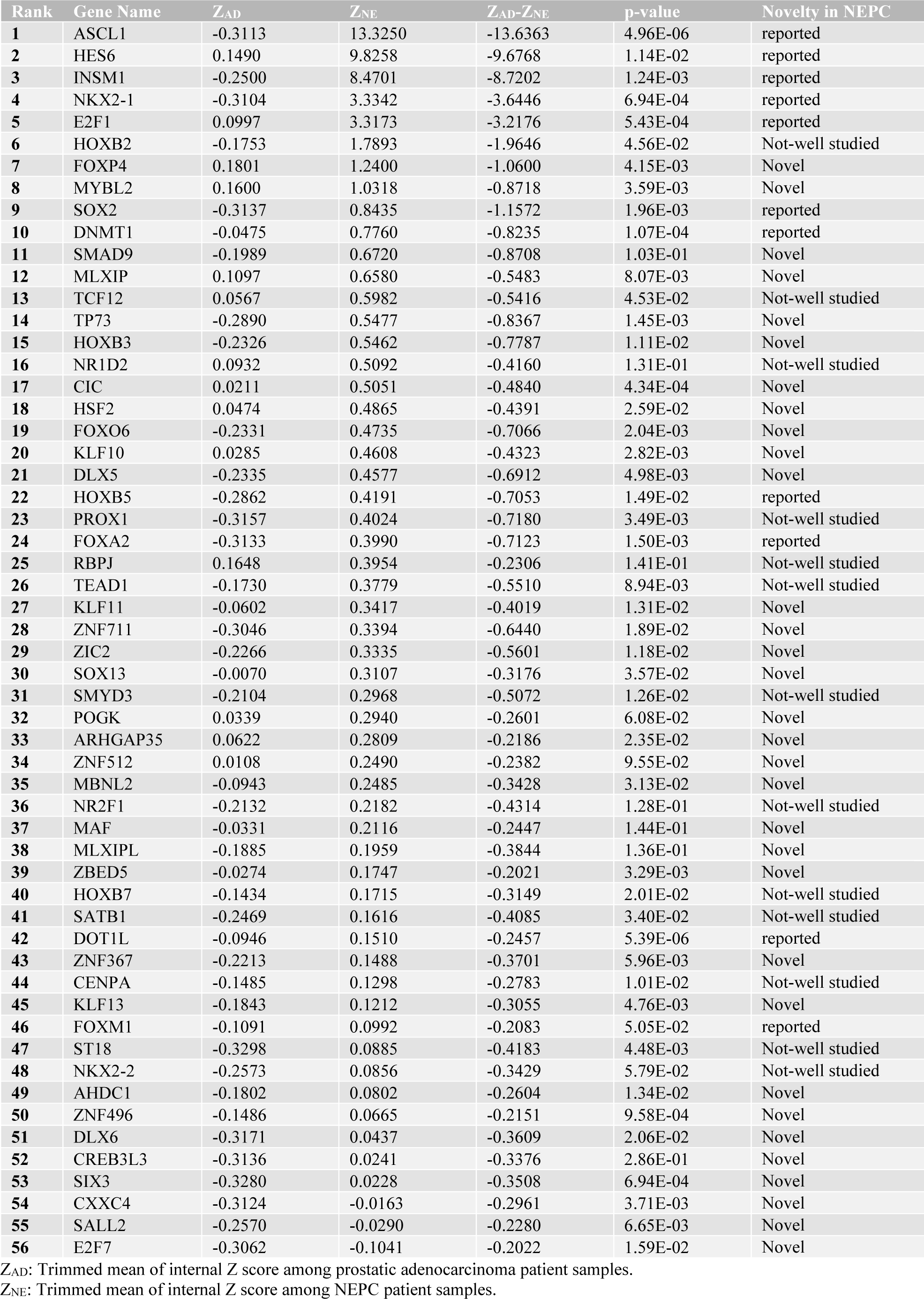
NE-TFs list.

### The lineage TF gene sets are clinically relevant and unveil novel candidates for PCa

To demonstrate the broad applicability of our TF gene sets in PCa, we employed gene set enrichment analysis (GSEA) across multiple PCa clinical cohorts. In Labrecque 2019 cohort (29), The AD-TFs gene set was enriched exclusively in adenocarcinoma, the NE-TFs gene set was enriched in NEPC samples, and the shared TFs gene set was enriched in both lineage (**Fig. 1D**). Unbiased hierarchical clustering distinctly separated the patient samples into two groups, underscoring the predictive capability of lineage-specific TFs (**Fig. 1, F and G**). This pattern was consistently observed in four additional clinical cohorts (**Fig. S2A**) and validated in two well-known PCa patient-derived xenograft (PDX) model series, the LTLs (12) and LuCAPs (30), demonstrating consistent lineage-specific TF enrichment **(Fig. S2, B, C and D**).

The comprehensive review of the lineage TFs gene sets revealed a substantial representation of well-characterized key TFs in both PRAD and NEPC (**Fig. 1E**). The AD-TFs gene set contained 21 previously identified TFs, such as NKX3-1 and MYC, with AR as the top ranking (**Fig. 1E, F**, **Table 1**). The NE-TFs set included 11 established TFs like ASCL1 and SOX2, plus several reported but not deepen-studied TFs indicating their potential as key regulators in NEPC (**Fig. 1G**, **Table 2**). The inclusion of these documented TFs further validates our TF profiles and introduces 20 novel AD-TFs and 32 novel NE-TFs (**Fig. 1E**, **Table 1** and **Table 2**), which hold promise as significant players or therapeutic targets for PRAD and NEPC. Notably, certain TFs exhibit specific high expression in PRAD compared to normal tissue (**Fig. S3**), such as ZNF761 and MESP1, linking these changes to poorer patient survival (**Fig. S4**), highlighting their clinical significance. Furthermore, examination of shared-high TF profiles reveals the presence of universally recognized oncogenic TFs such as YBX1, FOXA1, JUN, FOS, and STAT3 (**Fig. S5**), suggesting fundamental oncogenic features shared between the two PCa lineages.

### AD-TFs and NE-TFs functionally represent the corresponding lineage

Using bioinformatic analysis, we delved into the roles of lineage-specific TFs in cellular processes. Our findings revealed that shared TFs primarily oversee fundamental cell activities like signaling, metabolism, and survival, underscoring their universal role in basal cellular metabolism (**Fig. 2, A and D**). In contrast, TFs specific to adenocarcinoma or NEPC were more frequently involved in pathways governing cell development and differentiation, suggesting their potential role in lineage-directed processes (**Fig. 2, B and C**). Specifically, AD-TFs mainly affect the development of epithelial cell and gland (**Fig. 2G**), with key TFs listed in **Fig. 2H**. NE-TFs were linked to the development of neural or nervous system and the endocrine system (**Fig. 2G**), with key TFs listed in **Fig. 2I** and **Fig. 2J**. Notably, the AD-TFs enriched a pathway related to the negative regulation of neuron differentiation, suggesting potential roles of the involved TFs such like SOX9, REST, DLX1, THRB, and FOXO3 play function as suppressors of NE phenotype. In summary, our results show that AD-TFs and NE-TFs point to different developmental paths in PCa evolution.

**Fig. 2.**
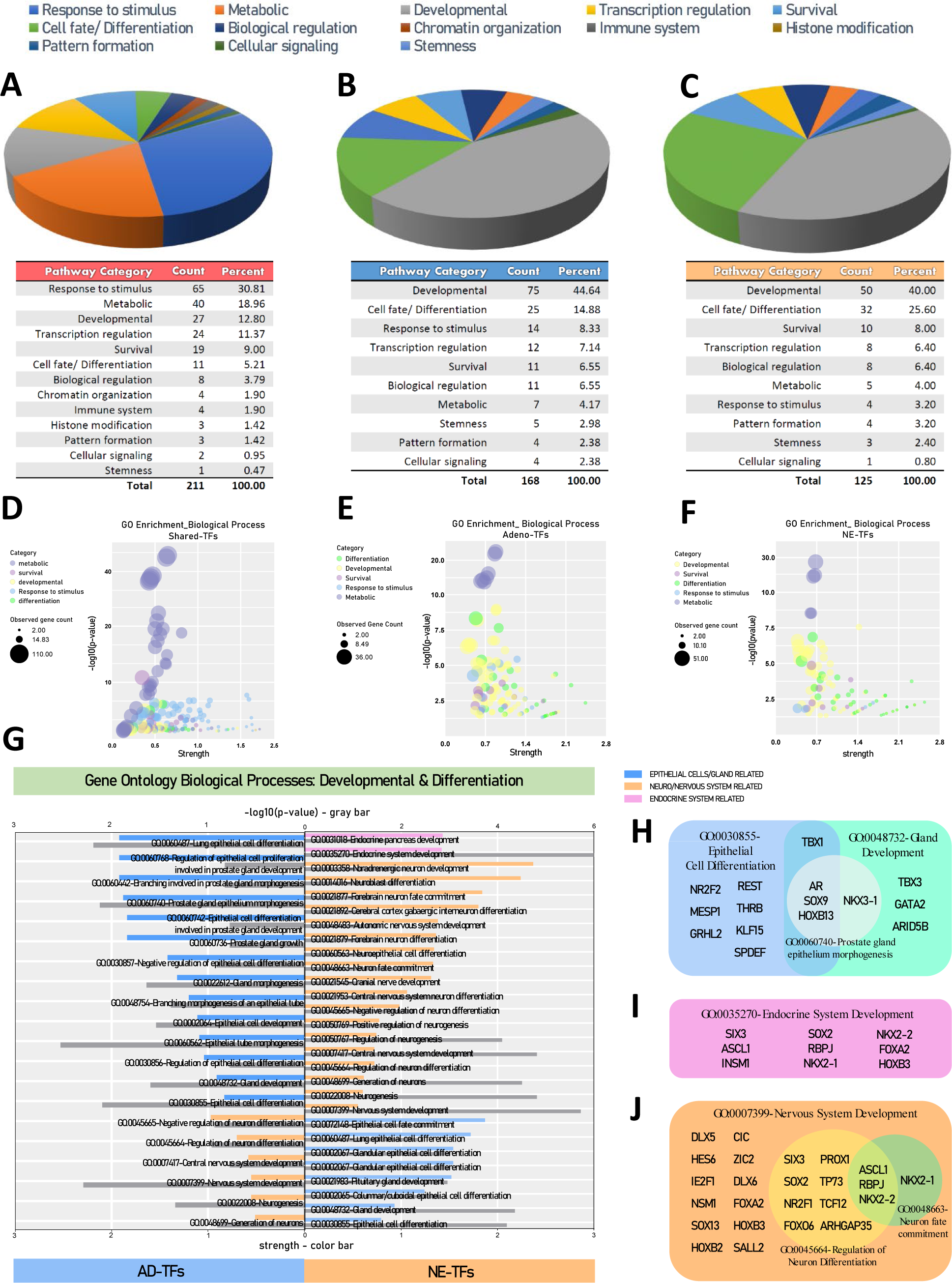
Functional Exploration of Lineage-TF Gene Sets. Pie charts illustrating the pathway distribution among (**A**) shared-TFs, (**B**) AD-TFs, and (**C**) NE-TFs, with detailed categories and proportions provided in the accompanying charts. Predominant pathway categories in shared-TFs include those related to stimulus response and metabolism, whereas over half of the enriched pathways in AD-TFs and NE-TFs pertain to developmental processes, cell fate and differentiation. Bubble plots (**D**) for shared-TFs, (**E**) AD-TFs, and (**F**) NE-TFs highlight pathway details for the top five categories (metabolic, survival, development, response to stimulus, and cell fate/differentiation), excluding the generic transcription regulation. (**G**) The enriched biological processes (Gene Ontology annotation) associated with developmental and differentiation in AD-TFs and NE-TFs exhibit divergent distribution in epithelial gland-related (Blue bar), neuro- and nervous-related (Orange bar), and endocrine system-related (Pink bar) pathways. All pathways shown have a p-value<0.01 (Gray-bar), with the lower x-axis indicating strength and the upper x-axis denoting -lg(p-value). Detailed TFs involved in key pathways of (**H**) epithelial cell/gland development-related, (**I**) endocrine system development-related, and (**J**) nervous system-related pathways are provided.

### Exploring novel lineage-specific TFs as potential therapeutic targets in PCa

According to the findings of the GO analysis, most lineage-specific TFs are concurrently implicated in metabolic-related processes, as reflected by the extremely low p-values and large gene counts (**Fig. 2, E** and **F**), underlining their pivotal role in cell survival and aggressiveness within their respective lineages. Drawing from the well-established roles of oncogenic TFs like AR and ASCL1 in PCa progression (17, 23), our investigation delved into the potential contributions of novel lineage-related TFs to cellular growth and aggressiveness. By identifying 10 novel AD-TF candidates (NFIX, THRB, ZNF613, ZNF614, ZNF615, CREB3L1, SIX5, ZBTB42, ZNF350, ZNF649) and 19 novel NE-TF candidates (AHDC1, CIC, CXXC4, DLX5, DLX6, DNMT1, HOXB3, HOXB7, HSF2, MAF, NKX2-2, NR1D2, RBPJ, SALL2, SATB1, ST18, TP73, ZNF367, ZNF711), we expanded the repertoire of potential therapeutic targets. Knockdown experiments targeting AD-TF candidates were conducted across a spectrum of androgen-response subtypes in PRAD cell lines, encompassing ADT-sensitive (LNCaP), ADT-insensitive (C4-2 and V16D), and ADT-resistant (22Rv1) adenocarcinomas. Results revealed varying degrees of inhibition in cell proliferation for most TF knockdowns (**Fig. 3, A, B, C and D**), with concurrent reduction in migration observed in V16D cells (**Fig. 3F**). In parallel, NE-TF candidates were subjected to knockdown experiments using 331R-2D, a NEPC cell line model derived from LTL331R PDX (31). Similarly, knockdown of NE-TF candidates led to inhibited cell proliferation (**Fig. 3E**). These findings underscore the pivotal role of lineage-specific TFs in tumor survival, suggesting their potential as promising therapeutic targets.

**Fig. 3.**
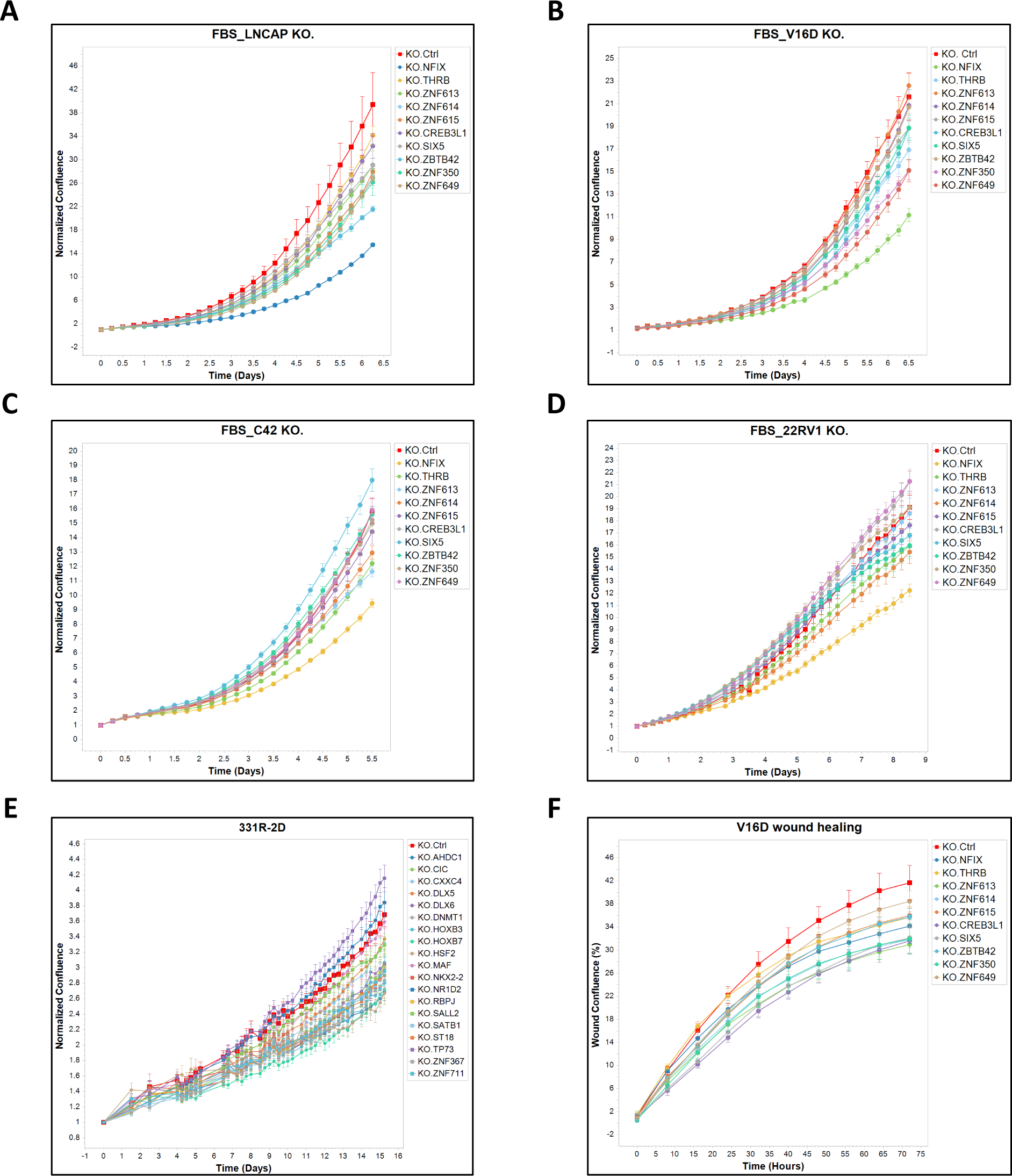
Impact of Lineage-TFs Depletion on Cell Growth and Migration in Different PCa Cell Lines. The proliferation of human PCa cell lines is compromised when infected with a lineage TF-targeted sgRNA construct, compared to a control CRISPR knockout lentivirus. Normalized confluence (relative to the initial point) was employed as a parameter to assess cell proliferation over time, as shown for (**A**) LNCaP cells, (**B**) V16D cells, (**C**) C4-2 cells, (**D**) 22Rv1 cells, which represent models of PRAD, and (**E**) 331R-2D cells, which is a cell model derived from NEPC PDX. (**F**) Wound confluence over time was used to evaluate the migration capability of V16D cells following AD-TF depletion. Data are represented as means ± standard deviation (SD).

### Deciphering the dynamics of Lineage TFs in NE transdifferentiation: A three-phase hypothesis

Our results revealed that lineage-TFs play pivotal roles in defining cellular lineage characteristics both molecularly and functionally, yet the transition of TF landscape from adenocarcinoma to neuroendocrine lineage is unclear. Utilizing the LTL331/331R transdifferentiation model, which recapitulates the development of NEPC from adenocarcinoma (12), we aimed to delineate the changes in lineage-TFs expression during this intricate process. Applying the internal z score algorithm, we observed structured changes in TF expression patterns during NE transdifferentiation: AD-TFs decrease significantly post-castration, persisting at low levels (**Fig. 4A**), while NE-TFs maintain low expression until relapse (**Fig. 4B**), suggesting a rapid loss of adenocarcinoma features upon ADT followed by a steep acquisition of NE features upon relapse. This reveals three discernible phases in the transdifferentiation process (**Fig. 4C**), informing our proposed three-phase hypothesis.

**Fig. 4.**
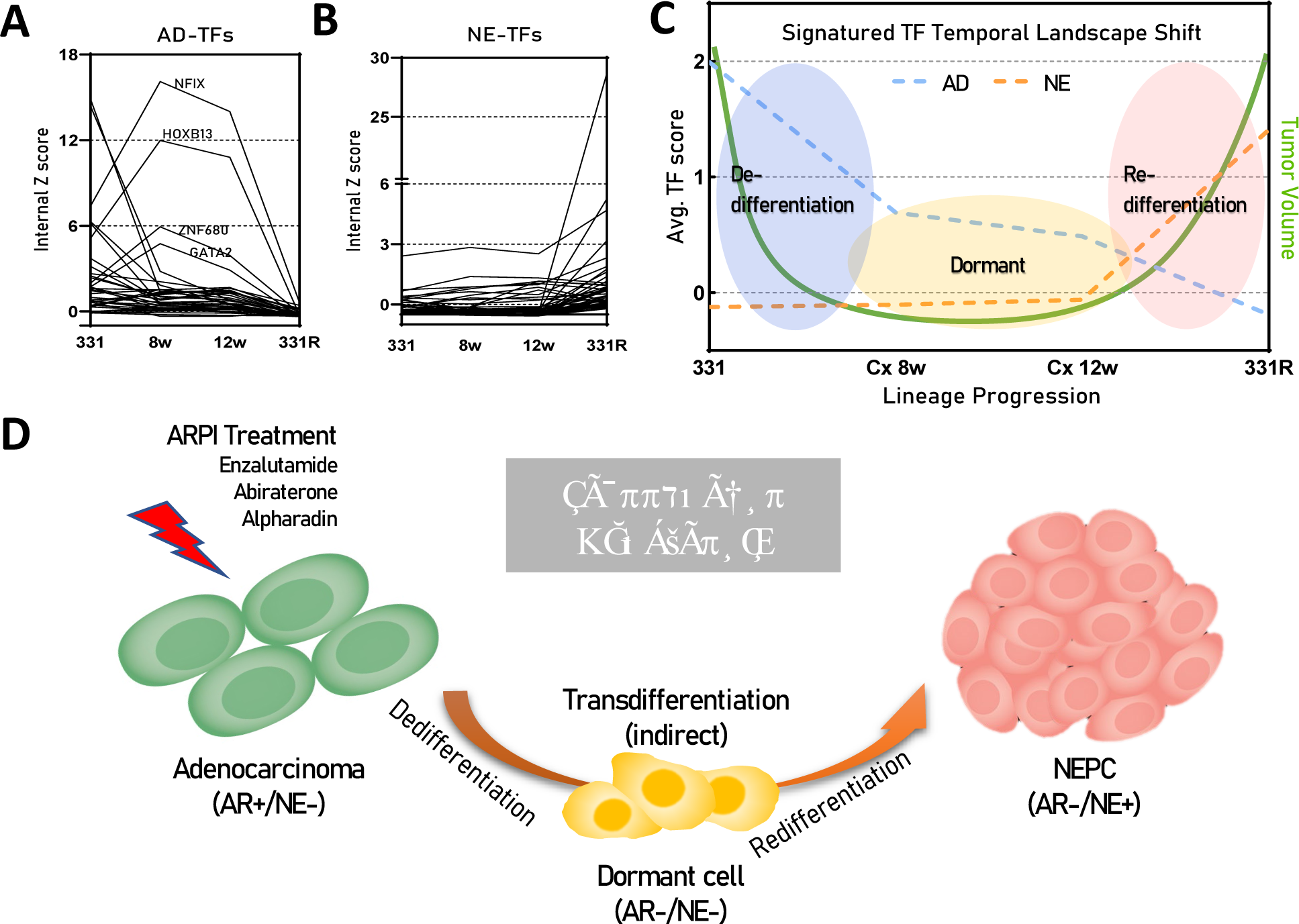
Proposal of the “Three-Phase Hypothesis”. (A) Internal Z scores of AD-TFs among the 331/331R time series model indicate reduced scores following castration of LTL331, with notable exceptions of NFIX, HOXB13, ZNF680, and GATA2 showing significant increases. *331: LTL331, PRAD; 8w: post-castrated 8 weeks of LTL331; 12w: post-castrated 12 weeks of LTL331; 331R: LTL331R, NEPC.* (B) Internal Z scores of NE-TFs among the 331/331R time series model show sustained low scores until the tumor relapse phase. (C) Trend of lineage-TFs shifts along the lineage progression timeline from LTL331 to LTL331R, represented by the average internal Z scores of AD-TFs (Blue dot line) and NE-TFs (Orange dot line), delineating the process into three distinct phases: de-differentiation, dormancy, and re-differentiation. Green line indicates tumor volume changes. (D) Graphic representation of the “Three-Phase Hypothesis”: AR+/NE-prostatic adenocarcinoma cells undergo de-differentiation under stress from AR pathway inhibition (e.g., enzalutamide, abiraterone, and apalutamide), entering a “dormant” state where both AD and NE features are suppressed. During a prolonged period of dormancy, cells acquire the capability for re-differentiation, eventually adopting the NE features to become NE+/AR-NEPC.

The first phase, termed the “de-differentiation phase,” marks the initiation of AD-TF loss upon ADT, indicating the adaptive survival of PCa cells capable of dormancy, distinct from androgen-dependent cells susceptible to ADT. Subsequently, dormancy-capable cells transition into a “dormancy phase,” marked by the repression of both AD-TF and NE-TF features, signifying an intermediate status. Following dormancy, cells may eventually acquire the capability to relapse, entering the “re-differentiation phase,” characterized by the acquisition of specific lineage features, leading to cancer relapse (**Fig. 4D**). Overall, our results provide a longitudinal analysis of lineage-specific TFs in NE transdifferentiation, offering a comprehensive view of the orchestrated molecular events that occur across three critical phases: de-differentiation, dormancy, and re-differentiation. These findings contribute to the elucidation of NEPC development and offer insights for potential therapeutic interventions targeting specific phases of this transdifferentiation process.

### TFs in dormancy: key roles in chromatin organization and maintenance of stemness

In the context of PCa, dormancy is a pivotal yet poorly understood phase in disease progression, largely due to the absence of appropriate models. Our hypothesis elucidated the dormancy phase in NE transdifferentiation as characterized by a sustained de-differentiated state with suppressed proliferation capability, pivotal for subsequent re-differentiation into NEPC. Notably, the transdifferentiation model LTL331/331R captures this dormant phase uniquely (12). The dormant state of LTL331/331R is characterized by a suppressed cell cycle capacity (**Fig. S6A**) and a reduced expression of the proliferation marker Ki67 (**Fig. S6B**), coupled with diminished AR / Prostate-Specific Antigen (PSA) expression indicative of an adenocarcinoma feature, and suppressed Chromogranin A (CHGA)/Cluster of Differentiation 56 (CD56) expression, which represent NE features (**Fig. S6B**). While most lineage-TFs conform to prevailing trends across the three phases, a minority, including NFIX, HOXB13, ZNF680, and GATA2, exhibit divergent patterns (**Fig. 4A**), suggesting the presence of additional TFs governing the dormancy phase.

Employing the internal Z score algorithm and screening criteria (**Fig. 5A**), we identified 213 TFs (**Fig. 5B**) with heightened weighted expression during dormancy compared to terminal stages (**Fig. 5C**). The protein-protein interaction (PPI) network analysis delineated dormant TFs into three distinct functional clusters (**Figure S7**). Cluster 1 TFs are significantly associated with histone modification and chromatin remodeling pathways (**Fig. 5, D and G**), indicating their role in transcriptional program restructuring during dormancy, with REST prominent within this cluster, a well-identified repressor of the NE phenotype, supporting that dormancy represents a repressed and de-differentiated state. TFs in cluster 2 are implicated in the regulation of stemness, governing pluripotency and stem cell population maintenance (**Fig. 5E**), indicating their potential role in preserving pluripotency for lineage plasticity. A few top-ranking transcription factors (TFs) within this group (**Fig. 5H**) have been previously associated with the initiation or preservation of dormant states across a variety of biological contexts such like STAT3 (32), FOXP1 (33), FOXO3 (34), SOX9 (35, 36) and SMADs family (37). In addition, the pathway enrichment of cytokine responses (**Fig. 5J**) implies potential intercellular communication via a ligand-receptor mechanism, such like the Interferon Regulatory Factor (IRF) family which regulate the Type I interferon system and adaptive immunity (38), suggesting their potential roles in protecting the tumor from immune surveillance.. Moreover, a substantial proportion of cluster 2 TFs are associated with pathways responsive to hormones (**Fig. 5I**). The upregulation of retinoic acid (RA) receptors (RAR/RXR genes: RARA, RARG, RXRA, RXRB) (**Fig. 5K**) provides insights into the potential involvement of RA in NEPC development, hinting at the potential mechanism underlying dormancy overcoming androgen receptor signaling abolishment. Overall, our findings shed light on the intricate TFs landscape of the dormancy phase during NE transdifferentiation, identifying key TFs and pathways that potentially govern this stage.

**Fig. 5.**
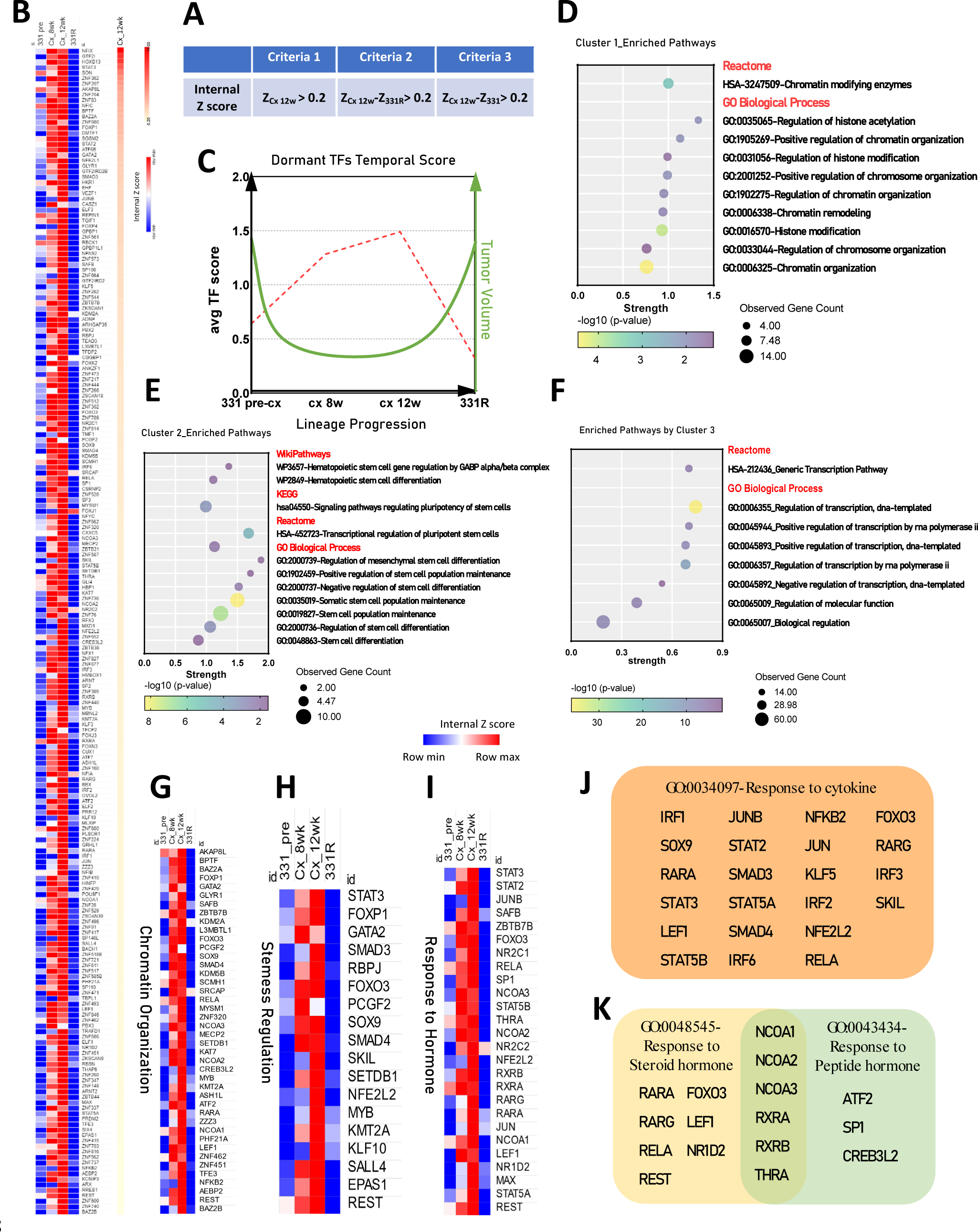
Exploration of dormant-TFs. (**A**) Criteria for screening of dormant TFs using the internal Z score. *Z_Cx 12w_: the TF internal Z score in post-castrated 12-week of LTL331. Z_331_/ Z_331R_: the TF internal Z score in LTL331/331R.* (**B**) Heatmaps displaying the signature of dormant-TFs in the LTL331/331R model, with TFs ranked based on Z_Cx 12w._ (**C**) Trend of dormant-TFs shifts along the lineage progression timeline from LTL331 to LTL331R, depicted by the average internal Z scores (Red dot line), alongside changes in tumor volume (Green line). (**D**) TFs in cluster 1 enriched pathways related to chromatin organization (**E**) TFs in cluster 2 enriched pathways associated with stemness regulation. (**F**) TFs in cluster 3 have enriched pathways related to generic transcription and biological processes. All pathways shown in bubble plots ing p-value<0.05. All pathways depicted in the bubble plots have a p-value < 0.05. Heatmaps detail expression patterns of key pathways enriched by dormant-TFs in (**G**) Chromatin Organization, (**H**) Stemness regulation, and (**I**) Response to hormone. Detailed TFs involved in certain pathways shown in (**J**) Response to cytokine, and (**K**) Response to hormone.

## Discussion

Our study pioneers a novel methodological approach to identify key regulators in PCa, specifically focusing on TFs involved in NE transdifferentiation. Unlike traditional analyses that prioritize DEGs, the internal Z score targets the weighted TF expression that enhances our ability to pinpoint pivotal regulators with subtle expression changes. This approach, scientifically robust and easily applicable, overcomes common barriers to practical use and holds promise for broad adoption across tumor types and diseases.

Our analysis reveals distinct transcriptional landscapes in PRAD and NEPC, emphasizing the role of TF profiles in cell lineage determination. By identifying lineage-specific TFs, we establish a molecular foundation for understanding adenocarcinoma to NEPC transitions, unveiling potential therapeutic targets. These verified lineage TFs provide novel candidates for further research.

Historically, the mechanisms of NE transdifferentiation have been poorly understood. Our work contributes to this area by illustrating the dynamic shifts in the lineage TFs landscape across three phases: de-differentiation, dormancy, and re-differentiation. We propose that the transition from adenocarcinoma to NEPC is an indirect process, challenging and expanding upon existing hypotheses like epithelial to mesenchymal transition (EMT) and epithelial immune cell-like transition (EIT).

The initial phase of de-differentiation, triggered by ADT, forces a small population of dormancy-capable cancer cells evading death by entering dormancy. Targeting this process could reduce dormant-capable cell populations, enhancing treatment efficacy. We’ve identified over 200 TFs related to chromatin remodeling and stemness during dormancy, offering potential targets to extend dormancy or achieve lifelong tumor suppression, aiming for non-recurrence.

In the re-differentiation phase, cells acquire terminal lineage features, often culminating in cancer relapses. Our findings suggest that intervention strategies should not only focus on treating NEPC but also on preventing progression to this final stage. By targeting early-stage vulnerabilities, particularly during the de-differentiation and dormancy phases, we can potentially control tumor development at its most treatable stages.

Overall, our study advances understanding of NE transdifferentiation in PCa, offering new research directions and potential therapies. By delineating TF interplay across cancer progression phases, we aim to shift from managing advanced disease to preventing its onset.

### Take Home Message

This study reveals lineage-related transcription factors in prostate cancer, highlighting their roles in cell development and providing novel potential therapeutic targets. It proposes a “three-phase hypothesis” for neuroendocrine transdifferentiation, offering insights into neuroendocrine prostate cancer progression.

## Material and Methods

### Patient-Derived Xenografts and Clinical Datasets

All LTL Patient-Derived Xenografts (PDX) lines were engrafted into NSG mice as previously described (12). This study followed the ethical guidelines stated in the Declaration of Helsinki, specimens were obtained from patients with their informed written consent form following a protocol (#H09-01628) approved by the Institutional Review Board of the University of British Columbia (UBC). All PDX microarray profiles are available at www.livingtumorlab.com and accessible under the accession number GSE41193 in the GEO database. The LTL331-castrated tissues and the NEPC-relapsed LTL331R tissues were harvested at different time points after host castration (39). Transcriptomic analysis of the LTL331/LTL331R time-series was conducted using RNA-sequencing data. RNA-seq profiles for the LTL331/331R castration time-series have been deposited to the European Nucleotide Archive and are available under the accession number ENA: PRJEB9660. The clinical cohorts utilized in this study include RNA-seq data from the following sources: Labrecque et al., 2019 (29), Abida et al., 2019 (40), Beltran et al., 2011 (41), Beltran et al., 2016 (42), Grasso et al., 2012 (43), and the LuCAP PDX series (30).

### TF score calculation and TF gene set identification

To calculate the weight expression for a TF, the following steps are performed, the algorithm schematic is shown in **Figure 1a**.

1. Obtain RNA-seq data matrix of the clinical cohort Labrecque et al., 2019 (29): this involves accessing the RNA-seq data for the clinical cohorts mentioned previously.
2. Extract all 1639 TFs: once the data matrix is obtained, extract the expression values for the 1639 transcription factors and obtain the TF data matrix. ***i*** represent the row number, ***a*** represent column number.

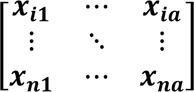
3. Calculate Column Average 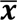 and Standard Deviation ***s*** : for each sample, calculate the average and standard deviation across all transcription factors.

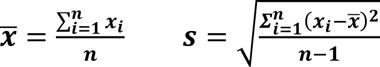
4. Internal Z-score calculation: calculate ***Z***_***i***_ for each TF in each sample and obtain the Z score data matrix.

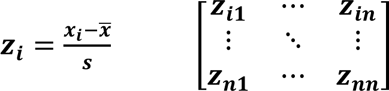
5. Calculate Z_ad_ and Z_NE_ using Trimmed Mean: for each TF, calculate the Trimmed Mean Z-score among either adenocarcinoma or NEPC samples, excluding the top and bottom 20% of Z-scores to reduce the impact of high-dispersion samples.
6. Adjustment of the internal Z-score threshold for grouping criteria: based on the expression levels of well-known transcription factors (TFs) in prostate adenocarcinoma and NEPC as reported in the literature, a threshold of 0.2 for the internal Z-score is considered appropriate for distinguishing TFs into relatively high-expressed or moderate/low-expressed categories.
7. TF gene sets classification by grouping criteria: the grouping criteria outlined in **Fig. 2B** were applied to categorize TFs into three distinct gene sets.
8. Repetition of algorithm for another clinical cohort: following the completion of TF gene set list generation for the initial cohort, the algorithm was repeated for an additional clinical cohort described by Abida et al., 2019 (40).
9. Overlap of TF Lists from two cohorts: to finalize the TF gene set lists, an overlap analysis was conducted between the TF lists obtained from the two cohorts. This analysis identified TFs common to both cohorts, thereby consolidating and refining the TF gene sets.

### Gene set enrichment analysis (GSEA)

GSEA (http://software.broadinstitute.org/gsea/index.jsp) was used in this study to determine whether a defined set of genes (e.g., AD-TFs, NE-TFs and shared-TFs) shows significant, concordant differences between two biological phenotypes (e.g., prostatic adenocarcinoma vs. NEPC). All GSEA analyses in this study used whole transcriptomic data without expression level cut-off as expression datasets. Reads counts were used for RNA-seq data from PDX models and clinical cohorts mentioned previously. A gene set was considered significantly enriched if its normalized enrichment score (NES) has an FDR q below 0.25.

### Gene Ontology (GO) enrichment analysis

GO enrichment analysis was performed to elucidate the biological processes associated with the TF gene sets identified in this study. TFs were annotated with GO terms using established databases and functional annotation tools (http://geneontology.org/). Enriched terms were visualized and interpreted to elucidate biological relevance. To facilitate interpretation, enriched GO terms were clustered into functional groups or categories using k-means clustering.

### Construction of Protein-Protein Interaction (PPI) network

The STRING database (https://string-db.org/) was utilized to construct a PPI network. The analysis was performed by submitting lists of TFs to the STRING database. Interactions among the submitted TFs were retrieved with pre-defined confidence thresholds to ensure high-quality interactions with various sources information integrated such as experimental data, computational prediction methods, and curated databases, to generate comprehensive interaction networks. Following the construction of the STRING PPI network, various network analysis techniques were applied to elucidate the functional and structural properties of the network. Measures such as node degree, and clustering coefficient were calculated to assess the topological characteristics of individual nodes within the network.

### Pan-cancer analysis

The online analysis platform TNMplot (https://tnmplot.com/analysis/) was utilized for a comprehensive pan-cancer analysis of gene expression in normal tissue compared within the paired tumor (44). The database of this tool comprises 56,938 samples, consisting of 33,520 samples from 3,180 gene chip-based studies, 11,010 samples from The Cancer Genome Atlas (TCGA) (including 394 metastatic, 9,886 tumorous, and 730 normal samples), 1,193 samples from Therapeutically Applicable Research to Generate Effective Treatments (TARGET) (comprising 1 metastatic, 1,180 tumorous, and 12 normal samples), and 11,215 normal samples from the Genotype-Tissue Expression (GTEx) project. Statistical significance was assessed using Mann–Whitney or Kruskal–Wallis tests. False Discovery Rate (FDR) was calculated using the Benjamini–Hochberg method.

### Overall survival analysis

cBioPortal (https://www.cbioportal.org/) was utilized to access all available PCa clinical cohorts. cBioPortal is an open-access resource that provides visualization, analysis, and download of large-scale cancer genomics datasets. We queried the cohorts for specific genetic alterations and visualized these using the OncoPrint feature. The Kaplan-Meier method was used to estimate survival functions, and the log-rank test was used to compare survival distributions between groups (e.g., altered vs. unaltered). All statistical analyses were performed using tools of cBioPortal.

### Cell lines and cell culture

The prostatic carcinoma cell lines LNCaP, C4-2, 22Rv1 and the human embryonic kidney cell line 293T cells were obtained from the American Type Culture Collection (ATCC; Rockville, USA). The castrate resistant prostatic carcinoma cell line V16D were derived through serial xenograft passage of LNCaP cells (45) which was a kind gift from Prof. Amina Zoubeidi laboratory (Vancouver Prostate Centre, Vancouver, Canada). The NEPC cell line 331R-2D were derived through serial xenograft passage of the LTL331R PDX model as previously described (31). Cells were authenticated with the fingerprinting method at Fred Hutchinson Cancer Research Centre (Seattle, USA). Mycoplasma testing was routinely performed at the Vancouver Prostate Centre (Vancouver, Canada). LNCaP, C4-2, V16D and 22Rv1 cells were maintained in RPMI-1640 medium (Gibco) containing 10% FBS (Gibco). 293T cells were kept in DMEM medium (Gibco) with 5% FBS. 331R-2D cells were cultured in RPMI-1640 medium with supplements as follows: 5% FBS, 10 nmol/L β-estradiol (Sigma-Aldrich), 10 nmol/L Hydrocortisone (Sigma-Aldrich), 10 µmol/L Y-27632 (Dihydrochloride, Stemcell), 1% Insulin-Transferrin-Selenium (Thermo Fisher) and 1% Matrigel (Corning).

### Vector Construction, Virus Production and Transduction

lentiCRISPR-v2 was a gift from Feng Zhang (Addgene plasmid #52961); pMD2.G was a gift from Didier Trono (Addgene plasmid #12259); psPAX2 was a gift from Didier Trono (Addgene plasmid #12260). Single-guide RNAs (sgRNAs) individually targeting 39 TFs were cloned into lentiCRISPR-v2 vector as previously described (46). Individual constructed lentiCRISPR-v2 vectors were co-transfected with psPAX2 and pMD2.G plasmids into 293T cells using Lipofectamine 3000 (Thermo Fisher) according to the manufacturer’s protocol. Culture media containing lentiviruses were collected at 48 and 72 hours respectively after virus packaging in 293T cells. Upon filtering, the virus-contained media were added to the culture media of either LNCaP, C4-2, V16D, 22Rv1 or 331R-2D cells, supplemented with 8 µg/mL polybrene (Sigma-Aldrich) to enhance transduction efficiency. Puromycin (Gibco) was used at 1 μg/ml to select for infected cells and maintain stable cell populations.

### Label-Free Cell Proliferation Assay

The Label-Free Cell Proliferation Assay was conducted using the IncuCyte® Live-Cell Analysis System. Cells were seeded into 96-well plates in complete medium at specified densities (**Table 3**), with three replicates per plate. The plates were then subjected to overnight incubation to facilitate cell attachment. Place the 96-wells plate inside the IncuCyte and allow the plate to warm to 37 °C for 30 minutes prior to scanning. Set the parameters for Scan type: Standard or Adherent Cell-by-Cell (for cell counting); Image channels: Phase; Objective: 10X; Scan interval: every 6 hours. Quantification of cell proliferation across multiple cell types is enabled using IncuCyte® AI Confluence Analysis provided within the IncuCyte™ software package.

**Table 3.** Cell seeding density for proliferation assay in 96-well plates.

| Cell line | LNCaP | C4-2 | V16D | 22Rv1 | 331R-2D |
| --- | --- | --- | --- | --- | --- |
| Seeding density (per well) | 2,500 | 2,000 | 2,000 | 4,000 | 20,000 |

### Migration analysis

The migration assay was conducted using the IncuCyte® live-cell imaging system with Scratch Wound Analysis Software Module. V16D cells were seeded into the 96-well ImageLock plate (Sartorius) at the density of 5,000/well with three replicates per plate, then allowed the cells grow to reach confluence. Wounds in all wells were simultaneously created using the 96-pin IncuCyte® WoundMaker according to the manufacturer’s protocol. After washing by PBS, complete medium was added into the plate which was then placed inside the IncuCyte. Set the parameters for Scan type: Scratch Wound; Image channels: Phase; Objective: 10X; Scan interval: every 8 hours. The data was then analyzed by the integrated metric of wound confluence using IncuCyte™ software package.

### Immunohistochemistry (IHC) analysis and antibodies

The harvested xenografts of LTL331/331R time series model were bisected along their longest dimension, fixed in 10% neutral-buffered formalin, and subsequently embedded in paraffin. Tissue sections were prepared from the formalin-fixed, paraffin-embedded samples, and routine hematoxylin and eosin (H&E) staining, as well as IHC, were performed following established protocols as previously described (12) using the primary antibodies including a rabbit monoclonal anti-AR antibody (1:100, Abcam), a rabbit monoclonal anti-PSA antibody (1:200, Abcam), a monoclonal mouse anti-human Ki-67 antibody (1:25, Dako), a rabbit monoclonal anti-CHGA antibody (1:500, Abcam) and a rabbit monoclonal anti-CD56 antibody (1:2000, Abcam).

### Statistical analysis and data representation

Statistical analysis was done using the Graphpad Prism software and the IncuCyte™ software package. The student t-test was used to analyze statistical significance between groups in discrete measurements, whereas two-way ANOVA was used for continuous measurements. The heatmap was constructed using Morpheus online tool (https://software.broadinstitute.org/morpheus/). The distance measure utilized in hierarchical clustering analysis was the Euclidean distance which served as the basis for clustering similar samples together in a hierarchical manner, facilitating the identification of underlying patterns and relationships within the dataset. The Kaplan–Meier method was used to estimate curves for relapse-free survival, and comparisons were made using the log-rank test. Any difference with P values lower than 0.05 is regarded as statistically significant, with *, P <0.05; **, P < 0.01; and ***, P < 0.001. Graphs show pooled data with error bars representing SEM obtained from at least three replicates and SD of values from clinical samples.

## Acknowledgments

We would like to thank all members of the Living Tumor Laboratory and collaborators for the thoughtful discussions.

## Author contributions

Conceptualization: YW, YZW

Methodology: YW, YZW

Xenograft studies: DL, HX, XD

H&E and IHC: RW

Advise on experiments: CK, FC

Data acquisition: CC, YL

Bioinformatic analysis: YW, YN, XZ, ZC, XP, *in vitro* experiment: YW, JC, MS, JC

Data interpreted: YW, XZ, YZW, AP, AC

Supervision: YZW, MEG, AC, CJO

Writing—original draft: YW, YZW, AP, AC

Writing—review & editing: YW, YZW, MEG, AC, CJO

## Data and materials availability

All data are available in the main text or the supplementary materials.

