## supplemental material for "Deciphering the Transcription Factor Landscape in Neuroendocrine Prostate Cancer Progression: A Novel Approach to Understand NE Transdifferentiation"

Yu Wang *et al.*

**This file includes:**

Figs. S1 to S7

**Fig. S1.**

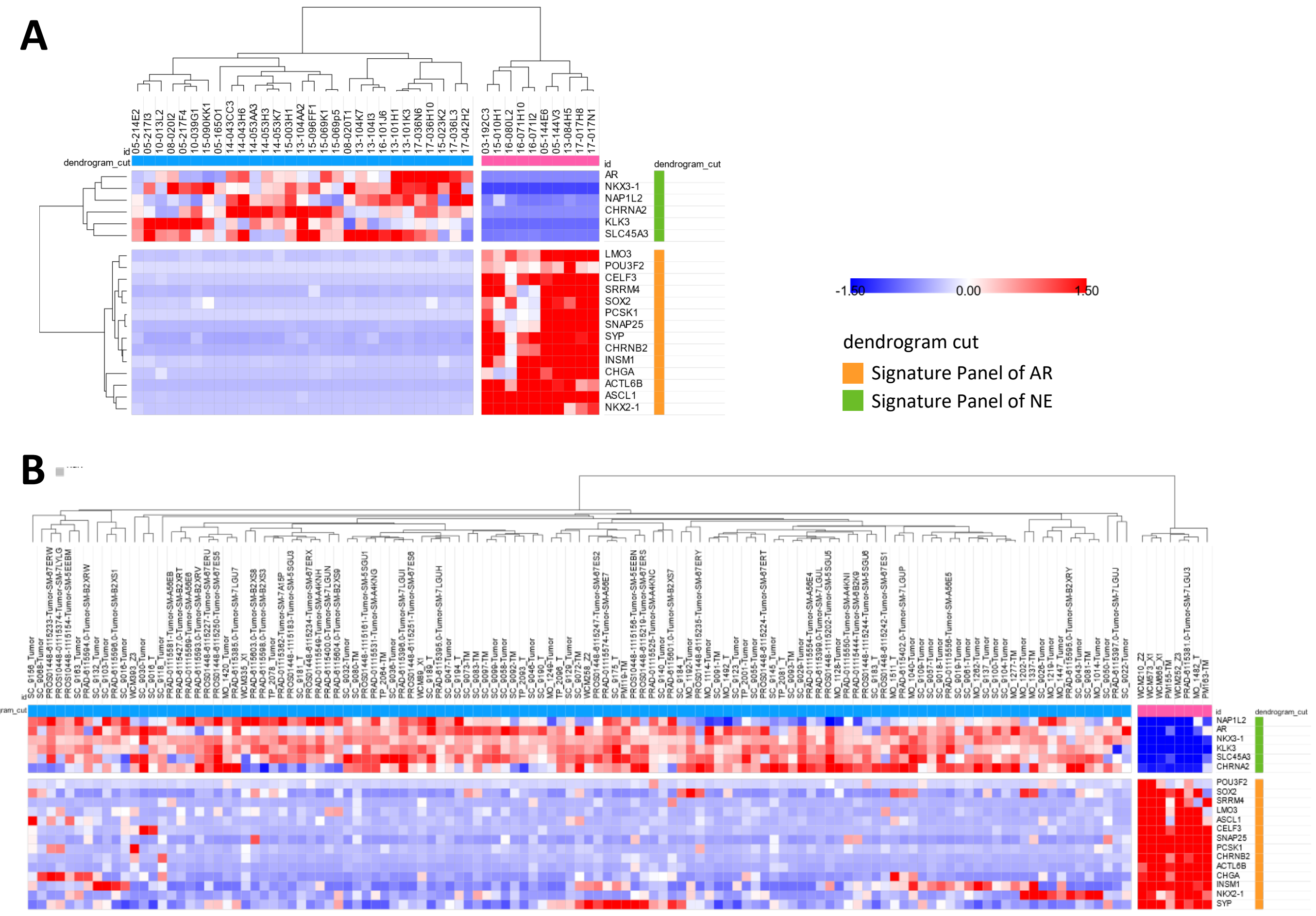


**Figure S1. Patient selection of the TFs-identification cohorts.** The heatmaps display patient samples for the identification of lineage-TFs in (**A**) the Labrecque et al. 2019 Cohort, and (**B**) the Abida et al. 2019 Cohort, using signature gene panels of AR and NE. Specifically, only patients classified as AR+/NE- for prostate adenocarcinoma (PRAD) and AR-/NE+ for neuroendocrine prostate cancer (NEPC) were included.

### Fig. S2.

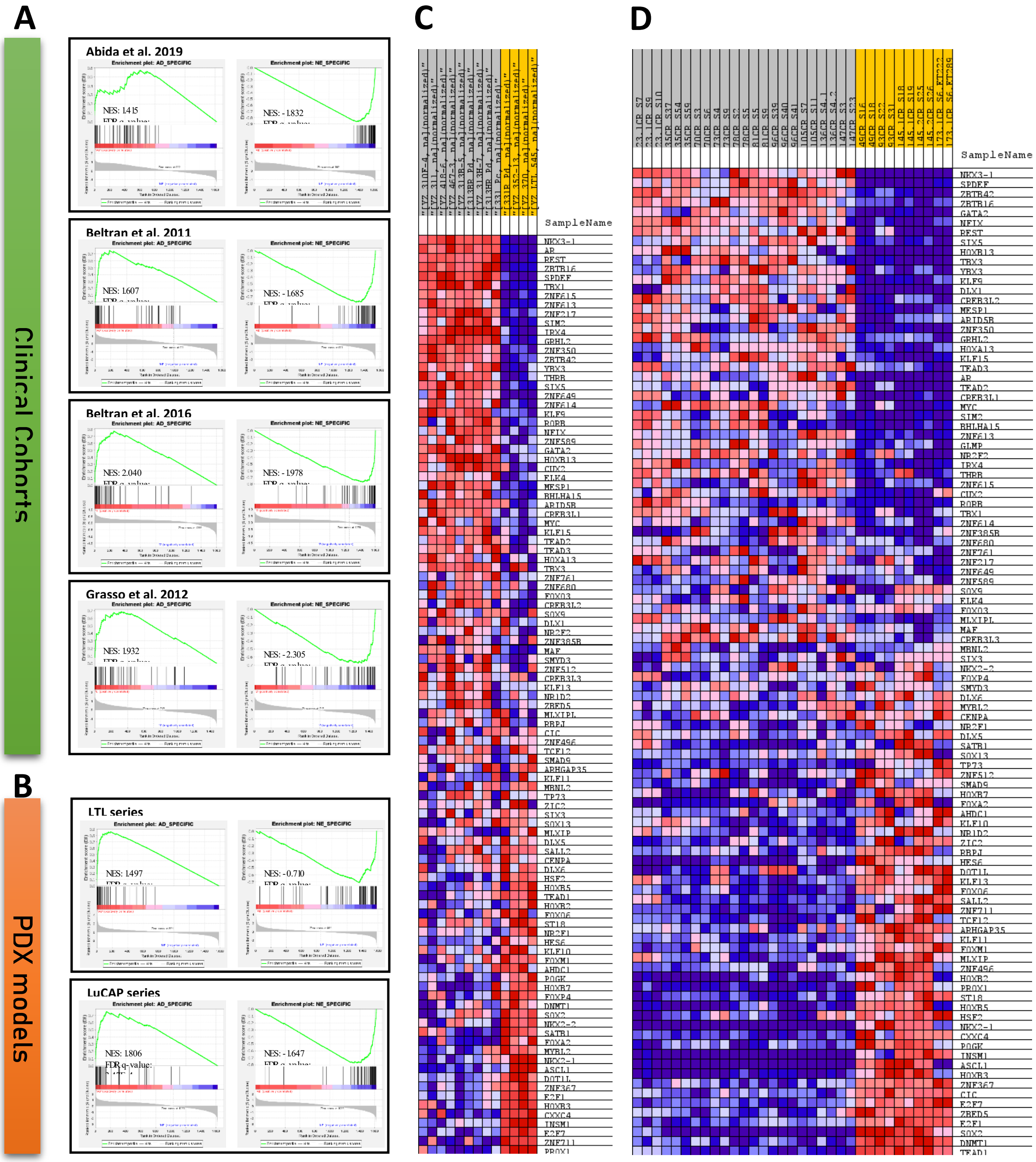


**Fig. S2. Validation of Lineage-TF Gene Sets in Clinical Cohorts and PCa PDX Models**. (**A**) GSEA validation of AD-TFs and NE-TFs gene sets in Abida et al. 2019 cohort (AD-TFs NES: 1.415, NE-TFs NES: -1.832), Beltran et al. 2011 cohort (AD-TFs NES: 1.607, NE-TFs NES: -1.685), Beltran et al. 2016 cohort (AD-TFs NES: 1.607, NE-TFs NES: -1.685), and Grasso et al. 2012 cohort (AD-TFs NES: 1.932, NE-TFs NES: -2.305). All cohorts show significant enrichment of lineage-TFs within respective patient lineages (FDR q-value<0.05). (**B**) GSEA enrichment of AD-TFs and NE-TFs in two series of well-established PCa patient-derived xenograft (PDX) models: LTL PDXs (AD-TFs NES: 1.497, NE-TFs NES: -0.710) and LuCAP PDXs (AD-TFs NES: 1.806, NE-TFs NES: -1.647), both with FDR q-value<0.05. Blue–Pink O' Gram in the Space of the Analyzed GeneSet representing the ratio values of all lineage-TF referred to 0 in the GeneSet of (**C**) LTL PDX models and (**D**) LuCAP PDX models. *NES: Normalized Enrichment Score. FDR: False Discovery Rate.*

### Figure. S3

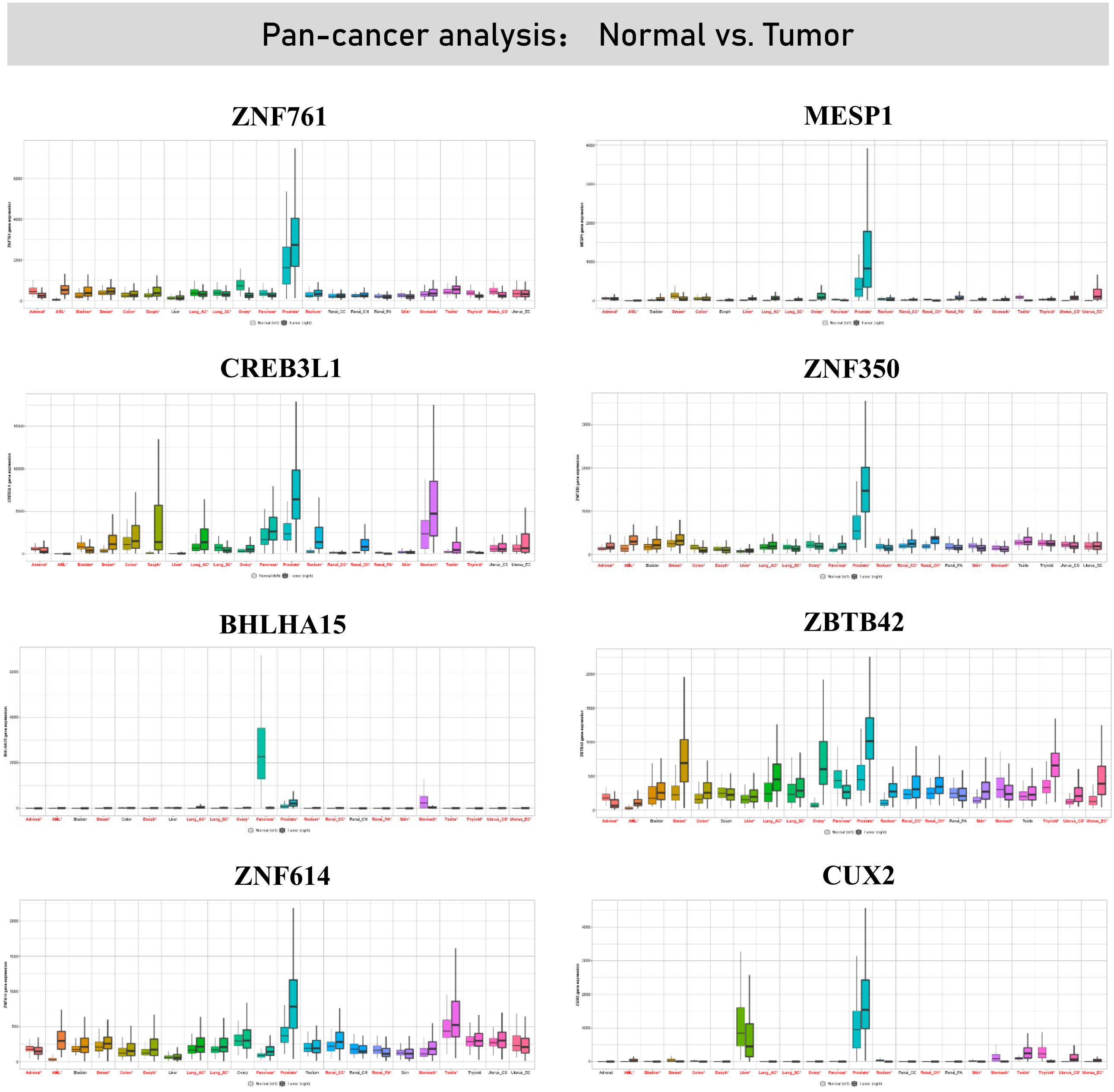


**Figure S3. Pan-cancer analysis of AD-TFs.** Expression Boxplots of AD-TFs including ZNF761, MESP1, CREB3L1, ZNF350, BHLHA15, ZBTB42, ZNF614, CUX2 in 22 common cancer types. The left box in each type of cancer indicates gene expression in corresponding normal tissue, the right box indicates the expression in tumor. Significant differences by a Mann–Whitney U test are marked with red color (* $p < 0.01$).

### Figure. S4

Prostatic adenocarcinoma

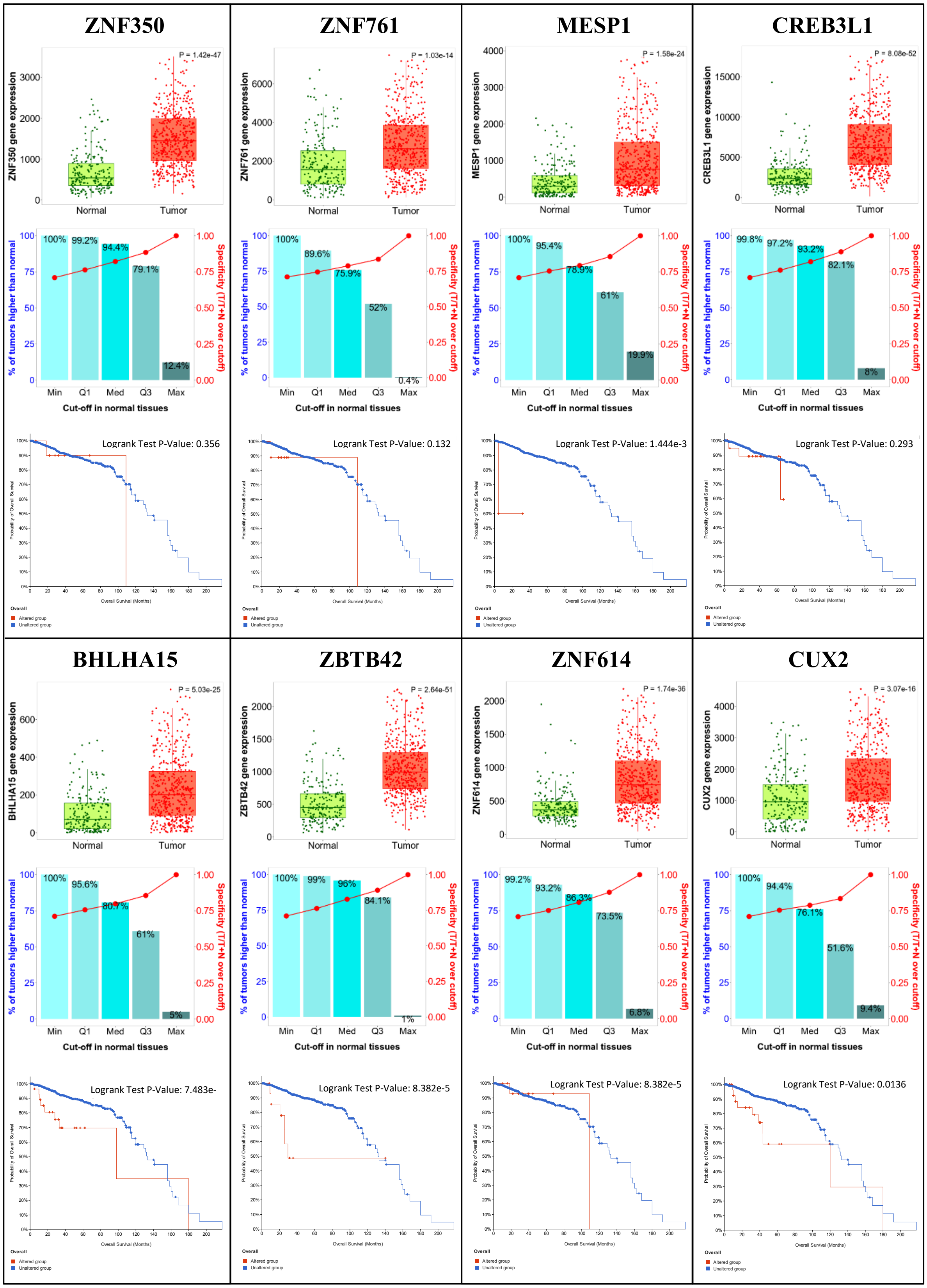

**Figure S4. The clinical relevance of AD-TF candidates in PRAD.** Boxplots indicate the gene expression in paired normal prostate tissue (Green box) or PRAD (Red box). The bars represent the proportions of tumor samples exhibiting higher expression of the selected gene compared to normal samples at each of the quantile cutoff values (minimum, 1st quartile, median, 3rd quartile, maximum). Specificity is calculated by dividing the number of tumor samples with values over each given cutoff by the sum of tumor and normal samples. For fold changes exceeding 1, those 'over' were considered instead of those 'below'. The Kaplan-Meier plot illustrates the overall survival of patients stratified by gene alteration status. The x-axis represents time (in months), and the y-axis represents the survival probability. The red line corresponds to patients with the gene alteration, while the blue line represents patients without the gene alteration. Differences in survival outcomes are statistically evaluated using the log-rank test.

### Figure. S5

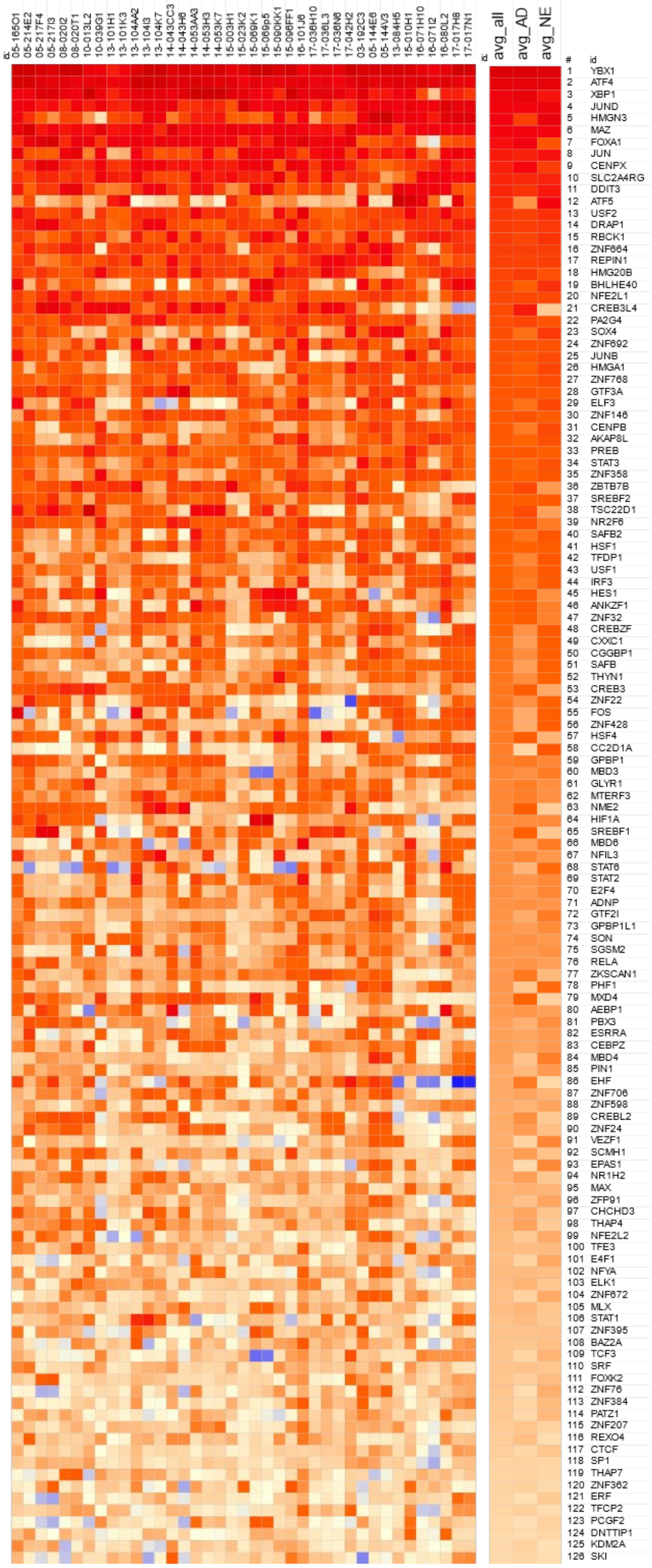


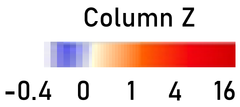


**Figure S5. Heatmaps exhibiting the signature of shared-TFs.** The heatmap visualizes the internal Z-scores of TFs shared by both PRAD and NEPC, ranking TFs based on the average internal Z-score across the two lineages.

#### Figure. S6

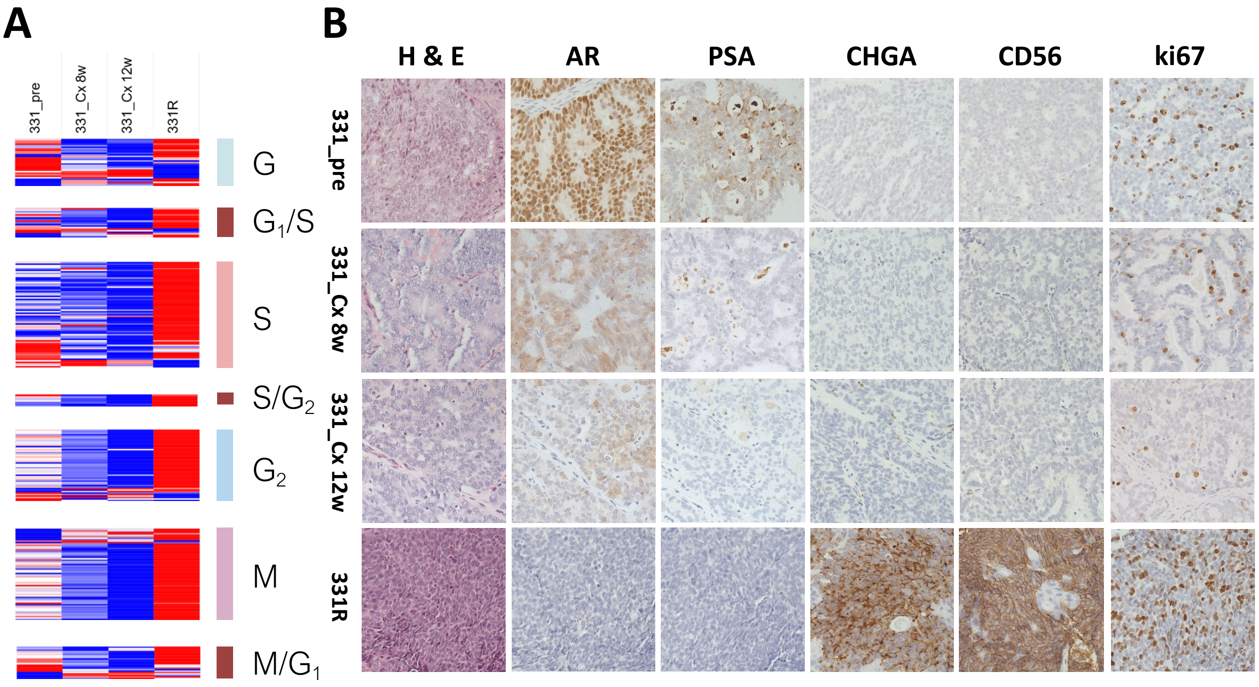


**Fig. S6. Dormant Features of the LTL331/331R Time Series Model.** (**A**) Expression Heatmap of cell cycle-related genes in LTL331/331R, with genes categorized by related cell cycle phase/checkpoint indicated on the right. (**B**) Immunohistochemical staining of AR pathway signal indicators (AR and prostate-specific antigen (PSA)), NE phenotype markers (chromogranin A (CHGA) and cluster of differentiation 56 (CD56/NCAM1)), and the proliferation marker Ki-67, alongside H&E staining depicting tissue original structure.

#### Figure. S7

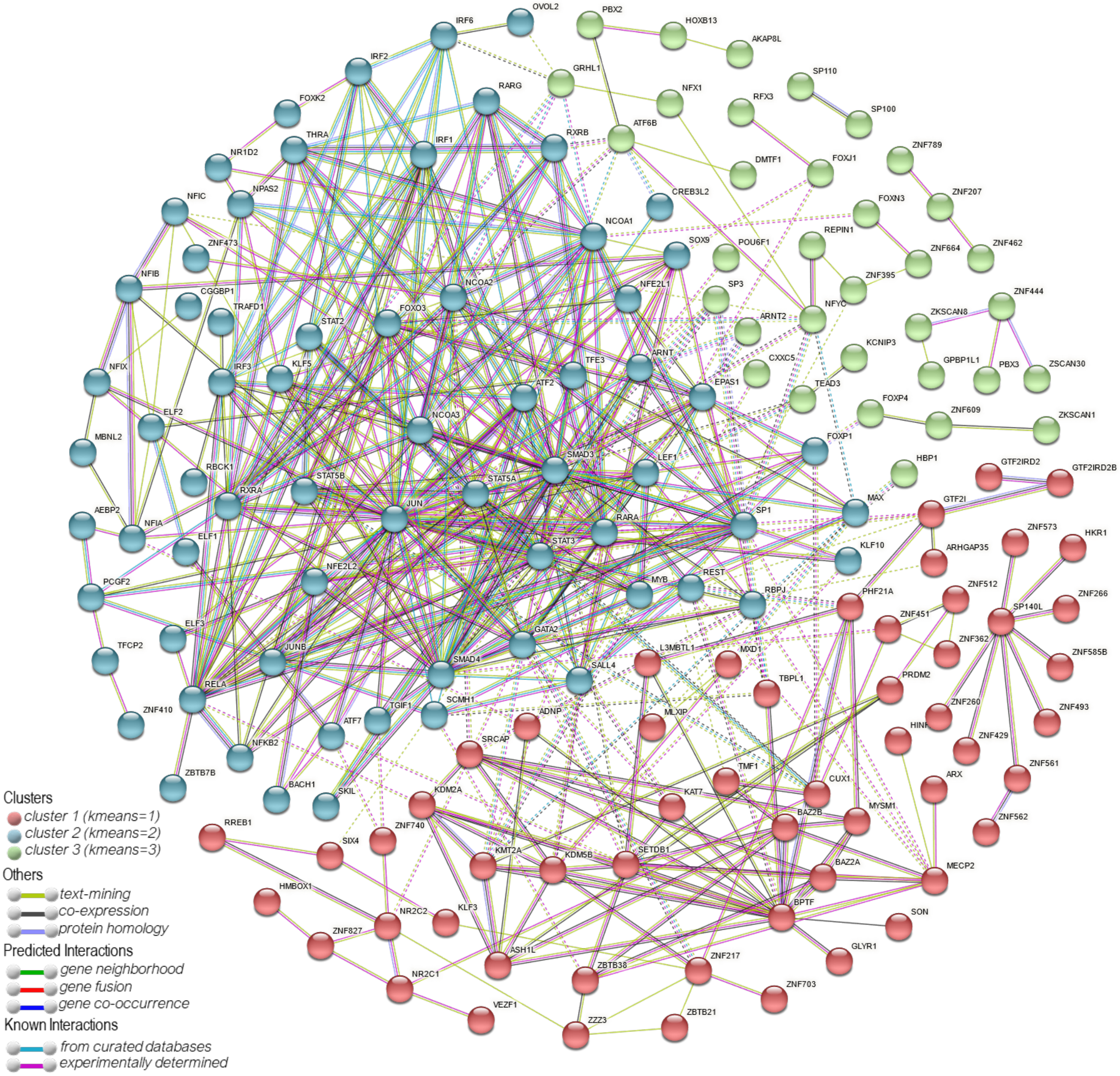


**Figure S7. The STRING protein-protein interaction (PPI) network among dormant-TFs.** The network diagram illustrates the protein-protein interactions (PPIs) among dormant TFs, identified through the STRING database. Each node in the network represents a TF, with edges between nodes indicating interactions between TFs as detailed in the legend. Clusters of densely interconnected nodes signify distinct protein complexes or functional modules.
